# Oral Intake of Deuterated Choline at Clinical Dose for Metabolic Imaging of Brain Tumors

**DOI:** 10.1101/2025.04.24.650452

**Authors:** Victor E. Osoliniec, Monique A. Thomas, Robin A. de Graaf, Henk M. De Feyter

**Author notes:** **Correspondence:** Henk M. De Feyter.

## Abstract

Accurate characterization and imaging of brain tumors are essential for effective treatment planning and monitoring. While MRI is widely used because of its high sensitivity for detecting lesions, the range of available types of MR image contrast does not offer high specificity for tumors. Deuterium metabolic imaging (DMI), which combines ^2^H magnetic resonance spectroscopic imaging (MRSI) with administration of deuterium-labeled substrates, is a relatively new imaging approach that could provide unique, complementary information to anatomical MRI. Preclinical studies have demonstrated the feasibility of DMI with intravenous (IV) administration of deuterated choline (^2^H_9_-Cho) for tumor characterization; however, they were performed at doses that exceeded severalfold the daily recommended Cho intake.

Here, we investigated the feasibility of oral (PO) administration of ^2^H_9_-Cho with a dose set at the recommended upper limit for daily use in humans. DMI was performed in rats with orthotopic glioblastoma tumors following a single, high-dose IV bolus (1 × 285 mg/kg) or low-dose PO administration over three consecutive days (3 × 50 mg/kg). Despite a lower cumulative dose, PO administration resulted in comparable total deuterated Cho (^2^H_9_-tCho) concentrations in the tumor, and tumor-to-brain image contrast relative to IV administration. Additionally, ^2^H and 2D ^1^H-^14^N HSQC NMR analyses on excised tumor tissue revealed differences in metabolite contributions to the in vivo ^2^H_9_-tCho peak. PO administration led to increased contributions from Cho-derived molecules that were products of tumor metabolism, than during IV infusion of ^2^H_9_-Cho. These findings suggest that repeated low-dose PO ^2^H_9_- Cho administration can generate high, image contrast between tumor and normal brain, that is predominantly generated by tumor metabolism instead of merely Cho uptake.

These results can advance the clinical translation of tCho-DMI as a noninvasive imaging tool for brain tumor characterization by demonstrating the feasibility of an oral intake approach using a clinically relevant dose. Given that Cho is already a widely used and well-tolerated nutritional supplement, oral Cho administration offers a practical, noninvasive alternative to IV infusion that could be conducted alongside regular MRI.

## Introduction

Non-invasive characterization of brain tumors remains a significant challenge. With its high sensitivity, anatomical magnetic resonance imaging (MRI) provides detailed structural information, but the low specificity of MRI cannot sufficiently characterize brain tumor lesions. This shortcoming becomes especially pronounced during treatment, when therapy itself can induce changes on structural MRI that are indistinguishable from brain tumor growth ^1–4^. Alternative methods, including existing metabolic imaging techniques, can offer unique complementary information to anatomical MRI ^5^. However, many of these techniques are limited by inconsistent tumor-to-normal-appearing brain (NAB) image contrast or challenges in clinical translation and underscore the need for more robust and reliable imaging approaches ^6^. One such novel method is deuterium metabolic imaging (DMI), a technique that integrates deuterium magnetic resonance spectroscopic imaging (^2^H MRSI) with the administration of a deuterated substrate of interest ^7,8^. Proof-of-concept experiments in a patient with glioma have already demonstrated that DMI-based maps, generated after administration of ^2^H-labeled glucose, showed image contrast with normal brain. This contrast originated from the aberrant glucose metabolism in high-grade brain tumors^7^. Another substrate with potentially high specificity for brain tumor imaging is choline (Cho), an essential nutrient involved in the synthesis of phospholipids, which are critical components of cell membranes ^9^. In many cancers, both Cho uptake and Cho metabolism are increased. Specifically, choline kinase alpha (CKA) is often upregulated in tumors, the first step in converting Cho to the downstream metabolites phosphocholine (PC) and glycerophosphocholine (GPC) in the Kennedy pathway (Fig. 1)^10–14^. The trimethyl groups of Cho, PC, and GPC can be routinely detected in vivo with ^1^H MRSI or ^2^H MRSI following deuterium labeling. However, for both methods, their similar chemical structures (Fig. 1) prevent spectral separation in vivo and result in a combined peak that is referred to as total Cho (tCho).

**Fig. 1:**
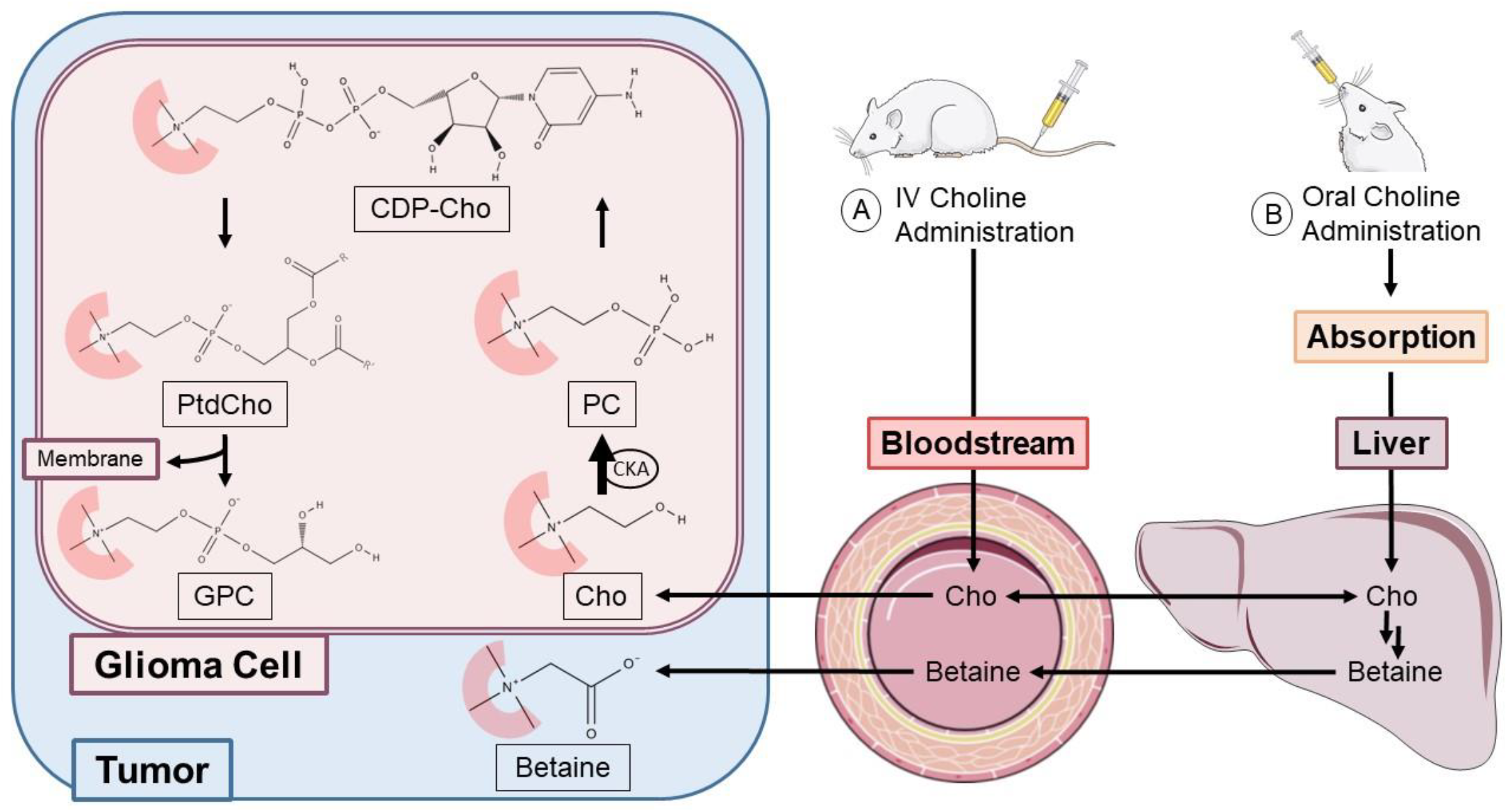
Overview of choline metabolism and study design. Schematic illustrating choline metabolism in glioma cells, and the differences between IV (A) and PO (B) administration of ^2^H_9_-choline. The ^2^H_9_ label, located on the trimethyl group, is highlighted in red on the molecular structures within the glioma cell. Previous studies have indicated increased choline kinase alpha (CKA) activity in cancer cells, thereby rapidly converting Cho compared to the slower conversion of PC to CDP-Cho. We hypothesize that this enzymatic rate difference can be leveraged with serial low PO doses of ^2^H_9_-Cho to accumulate a ^2^H_9_-PC pool that results in tumor-to-NAB contrast comparable to a single, high IV dose. PO administration implies that ^2^H_9_- Cho enters the circulation via the liver, where betaine is synthesized from Cho. Betaine could contribute to the DMI maps, however, it has not been shown to be related to tumor metabolism, and it remains unclear whether betaine is taken up by RG2 glioma cells or remains extracellular. Abbreviations: Cho: choline, PC: phosphocholine, CDP-Cho: cytidine diphosphate-choline, PtdCho: phosphatidylcholine, GPC: glycerophosphocholine, CKA: choline kinase alpha. Parts of the figure were adapted from Servier Medical Art, licensed under a Creative Commons Attribution 4.0 Unported License (https://creativecommons.org/licenses/by/4.0/).

Previous research demonstrated that tCho-based DMI can provide a high signal-to-noise ratio (SNR) for deuterated tCho and generate high tumor-to-NAB image contrast in a rodent model of glioblastoma (GBM), both *during* and *after* intravenous (IV) infusion of deuterated choline (^2^H_9_-Cho). Twenty-four hours after a single IV dose of ^2^H_9_-Cho, deuterated tCho in tumors was only ~25% lower compared to the levels detected in vivo during infusion ^15^. Furthermore, using high-resolution ^2^H nuclear magnetic resonance (NMR) on metabolite extracts from excised tumor tissue, it was shown that the residual ^2^H_9_-tCho signal consisted mostly of ^2^H-labeled PC and GPC. These previous results indicated a higher uptake and conversion rate of free Cho to PC compared to the slower metabolism downstream of PC (Fig. 1) ^16–18^.

In translating tCho-DMI to humans, the potential side effects of IV Cho infusion could represent a significant hurdle. Though most of the possible side effects can be considered relatively mild, high plasma levels of Cho after IV infusion can lead to lower blood pressure via cholinergic stimulation, which raises safety concerns for potential clinical use ^19^. Alternatively, oral intake of Cho is safe and often used as a nutritional supplement, with 3.5 g recommended as the maximum daily intake for adults ^20^. We hypothesized that the natural metabolic rate differences within the Cho pathway could be leveraged to accumulate ^2^H_9_-tCho signal in vivo with repeated oral doses that are at the maximum recommended daily intake for adults, and combined >5 times lower than the amount of Cho used so far in previous IV protocols. Specifically, by administering ^2^H_9_-Cho over multiple consecutive days, at an equivalent dose recommended for humans, we anticipated that the ^2^H_9_-tCho peak detected in vivo would increase with each dose. We also predicted that the ^2^H_9_-tCho peak would consist mostly of ^2^H-labeled PC and GPC, as with every oral dose, free ^2^H_9_-Cho would rapidly enter the tumor and get phosphorylated to PC. Further conversion of labeled PC is relatively slow, while the remaining ^2^H_9_-Cho in plasma is rapidly cleared ^21,22^. Each consecutive oral dose could therefore add to the existing PC labeling and increase the in vivo ^2^H_9_- tCho peak.

This strategy was tested in a rat model of GBM by measuring the [^2^H_9_-tCho] of tumors and NAB, in vivo with DMI. Tumor-to-NAB image contrast was compared between animals receiving a single, high IV dose and animals that were administered multiple low doses PO. To identify the different Cho metabolites that contribute to the in vivo ^2^H_9_-tCho peak, tumor tissue was harvested at the end of the in vivo experiments. ^2^H NMR and 2D ^1^H-^14^N Heteronuclear Single Quantum Coherence (HSQC) NMR experiments were performed on metabolite extracts from excised tumor tissue to detect ^2^H-labeled Cho, PC and GPC, as well as betaine, a metabolite formed in liver and kidneys that could also contribute to the ^2^H_9_-tCho peak in vivo but would not be specific for tumor metabolism ^23^.

## Results

### In vivo

All tumor-bearing rats showed a clear lesion on contrast-enhanced T_1_-weighted MRI (CE T_1_W MRI). This confirmed that the blood-brain barrier was compromised, a characteristic of GBM and congruent with previous reports using the RG2 model (Figure 2A). Anatomical MRIs were used to create segmentations of the tumor and NAB. For the IV group, the tumor volume-of-interest (VOI) range was 38.2 to 187.5 mm^3^ (mean ± SD: 103 ± 51 mm^3^), and the range of NAB values was 52.7 to 188.4 mm^3^ (mean ± SD: 107 ± 40 mm^3^). In the PO group, the range of tumor volumes was 62.5 to 187.9 mm^3^ (mean ± SD: 136 ± 39 mm^3^), and 54.7 to 175.0 mm^3^ (mean ± SD: 134 ± 39 mm^3^) for NAB volumes.

**Fig. 2:**
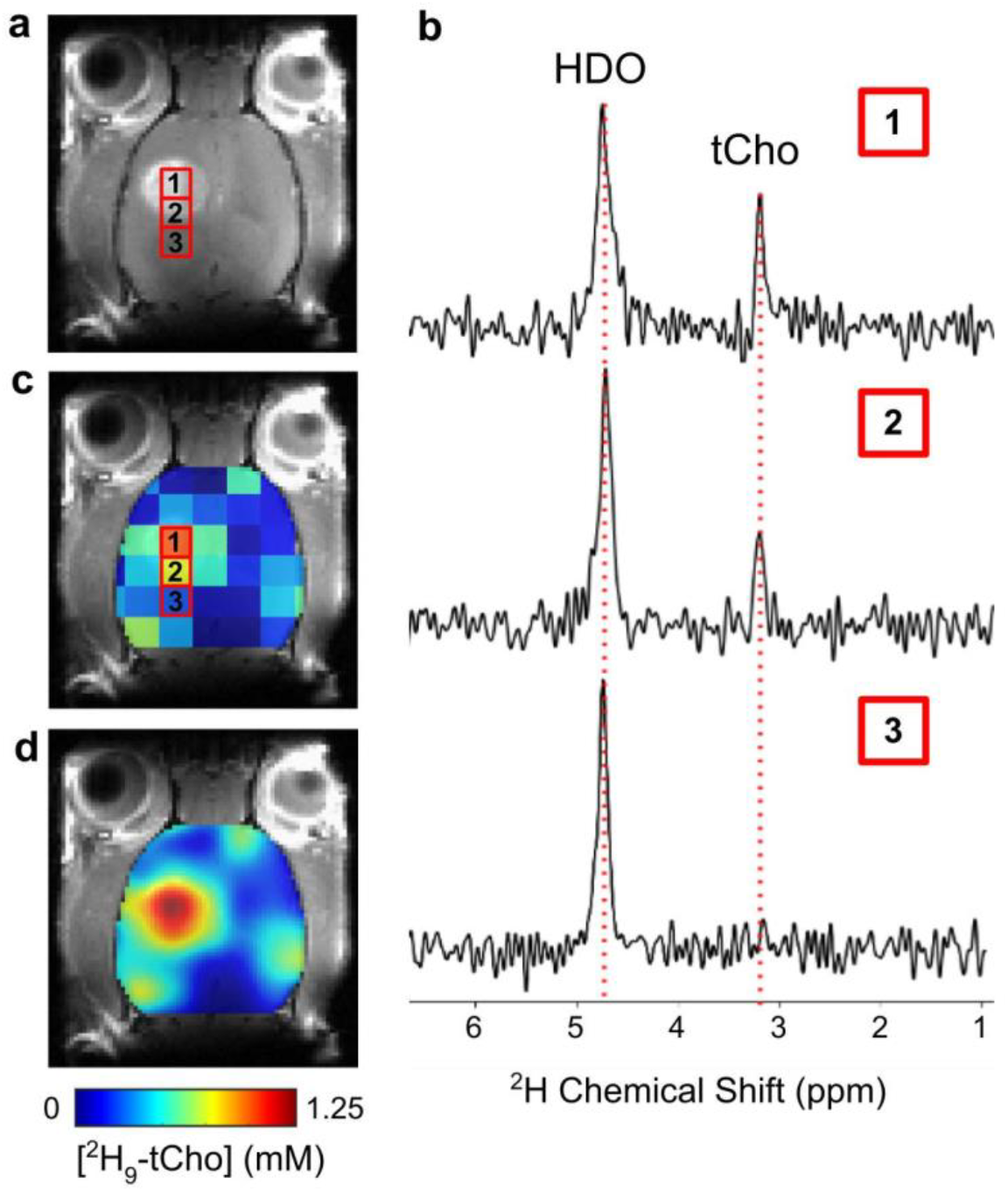
DMI following administration of ^2^H_9_-Cho. **a** Coronal slice of CE T_1_W MRI from an RG2-bearing rat, showing the tumor lesion and individual voxel positions selected from the ^2^H MRSI grid. **b** Individual ^2^H spectra (phase-corrected and line broadened with a 3 Hz Lorentzian function) from voxel positions indicated on the MRI shown in panel **a. c** Color-coded ^2^H MRSI grid overlaid on anatomical MRI. **d** Interpolated color-coded map based on data shown in panel **c**. Color scale applies to panels **c** and **d**. tCho: total choline; HDO: ^2^H-labeled H_2_O.

Figure 2 shows an example of the raw data obtained from an animal after three consecutive days of ^2^H_9_-Cho administration PO. The figure panels show ^2^H MRSI spectra, color-coded values of peak fitting results, and smoothed metabolic maps of [^2^H_9_-tCho] overlaid on anatomical MRIs. Assessment of individual spectra shows that voxels containing a mix of tumor and NAB have a lower [^2^H_9_-tCho] than a voxel located entirely within the contrast-enhancing lesion, which contains almost exclusively tumor tissue.

To test if [^2^H_9_-tCho] was increasing with each additional gavage dose, two animals underwent DMI scans on each day of PO administration. The average tumor [^2^H_9_-tCho] was near the noise level on day 1, increasing to 0.51 ± 0.02 mM on day 2 and 0.66 ± 0.15 mM on day 3 (n = 2). The corresponding metabolic maps reflect this steady increase in tumor [^2^H_9_-tCho], as shown in Figure 3.

**Fig. 3:**
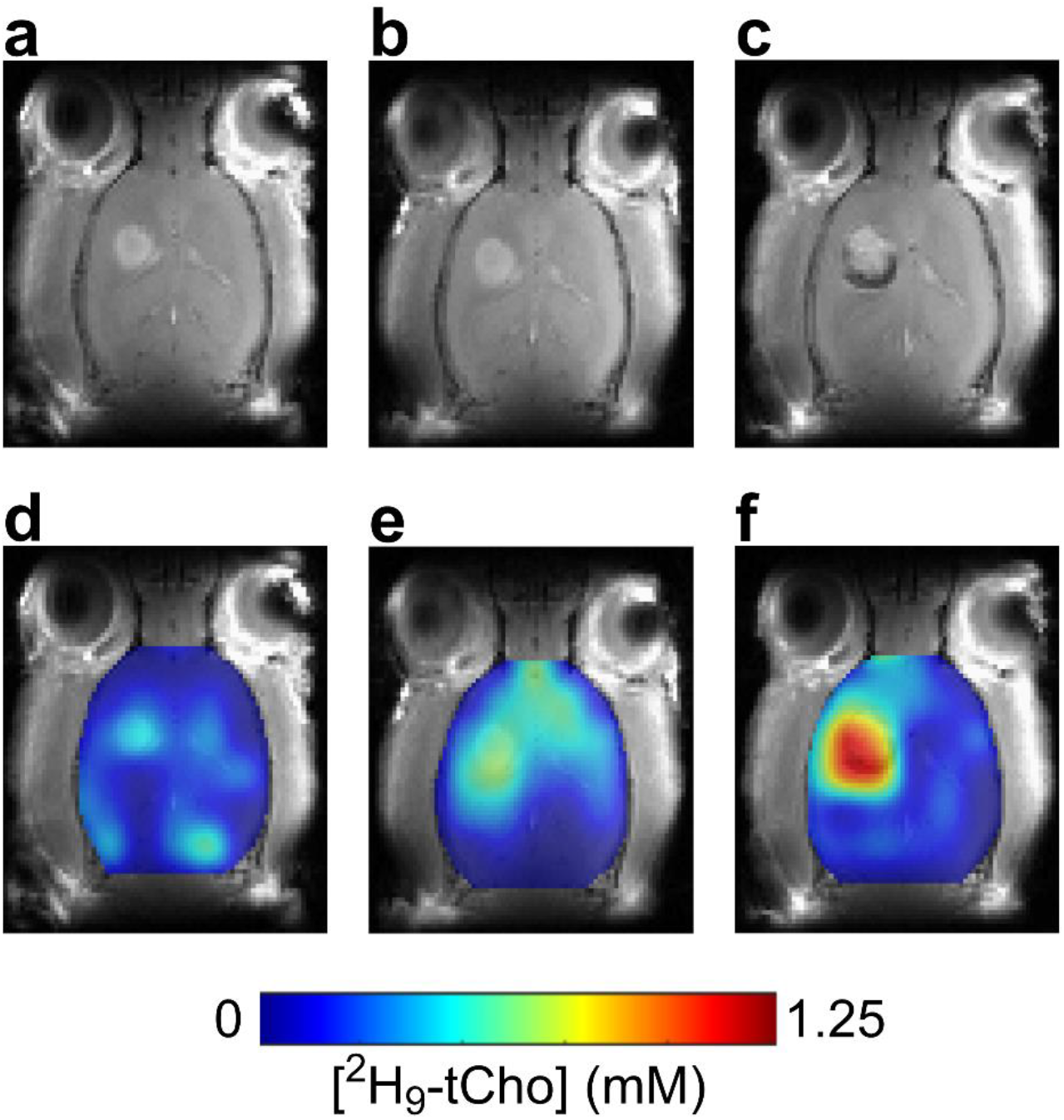
Buildup of tCho-DMI image contrast following serial oral doses. Example of a tumor-bearing rat that was scanned for each day of PO ^2^H_9_- Cho administration. CE T1W MRIs (**a – c**), and accompanying DMI-based maps of ^2^H_9_-tCho concentration (mM) (**d – f**).

Visual comparison of metabolic maps showed minimal differences between the groups of IV and 3 days of PO ^2^H_9_-Cho administration, as shown by representative data in Figure 4. To measure tissue-specific [^2^H_9_-tCho], VOIs of tumor and NAB were multiplied with the tCho-DMI-based metabolic maps (Fig. S2, Fig. 5). The average [^2^H_9_-tCho] following IV administration (n = 10) was 0.67 ± 0.15 mM in tumors and 0.23 ± 0.08 mM for NAB. The [^2^H_9_-tCho] in animals that underwent 3 days of PO administration (n = 9) was 0.70 ± 0.22 mM in tumors (p = 0.62 vs. IV group) and 0.20 ± 0.06 mM in NAB (p = 0.35 vs. IV group). The average tumor-to-NAB [^2^H_9_-tCho] ratio was 3.3 ± 1.6 for the IV group and 3.9 ± 1.7 for the PO group (p = 0.28) (Fig. 5).

**Fig. 4:**
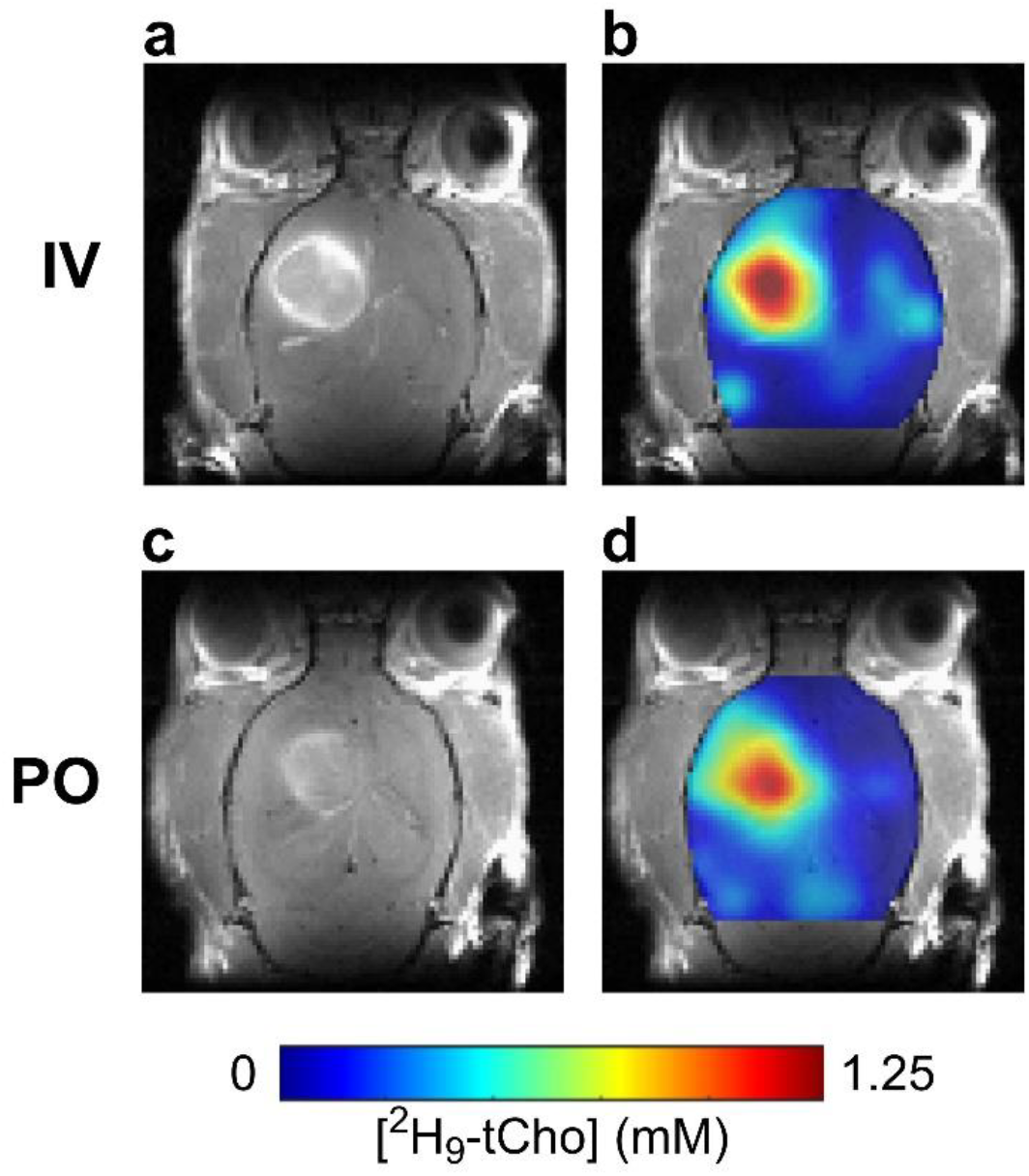
Comparison of in vivo DMI maps between IV and PO administration. **a, c** CE T_1_W MRI of RG2-bearing rats acquired during DMI scan sessions. **b, d** color-coded maps of [^2^H_9_-tCho] based on DMI data acquired during 36 min IV infusion of ^2^H_9_-Cho (**b**), and ~ 2 hours after PO gavage of ^2^H_9_-Cho on day 3 (**d**).

**Fig. 5:**
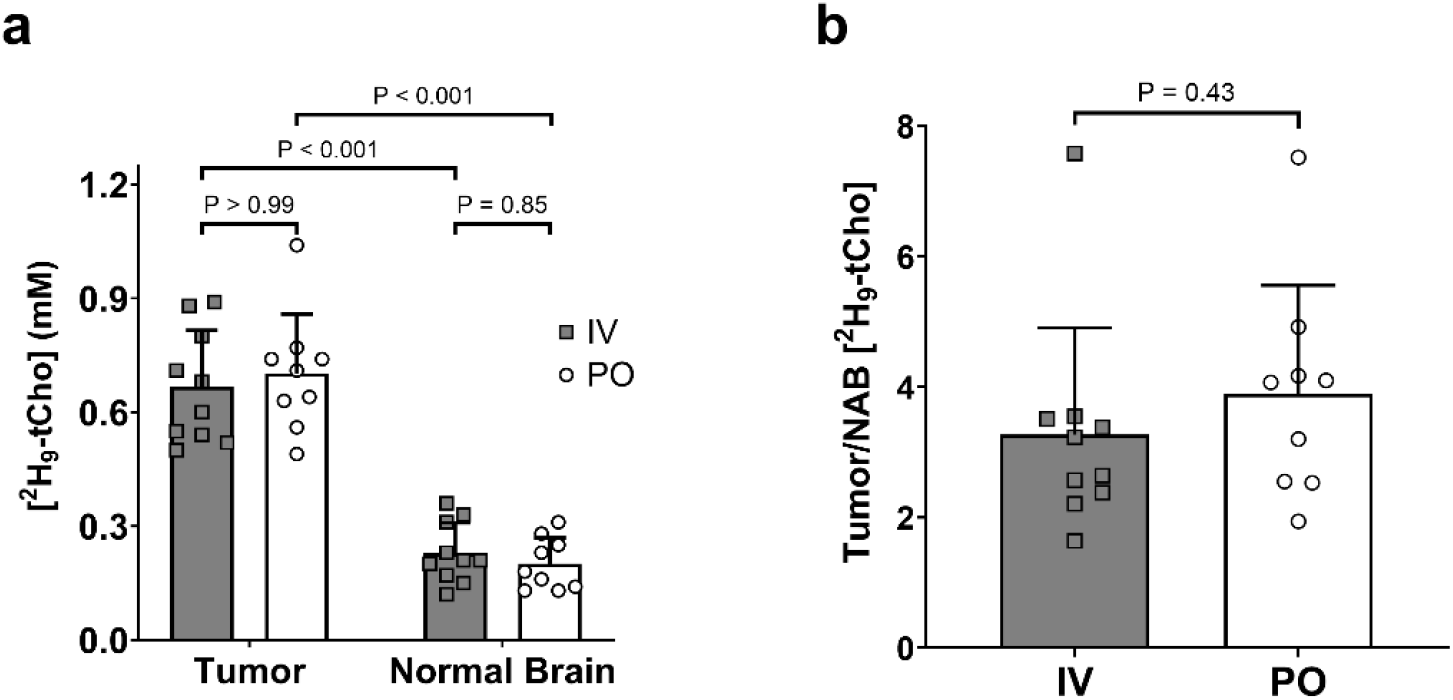
Quantification of in vivo [^2^H_9_-tCho]. **a** [^2^H_9_-tCho] (mM) in tumor and NAB for IV (n = 10) and PO (n = 9) administration groups. **b** Tumor-to-NAB image contrast for IV and PO administration groups. All plots indicate individual animal values and mean ± SD error bars. P values represent the results of two-tailed, independent sample t-test.

### Ex vivo

To identify the different Cho species that contribute to the single ^2^H_9_-tCho peak observed in vivo, high resolution NMR was performed on metabolite extracts generated from excised tumor tissue (Fig. 6). Analysis of the ^2^H NMR data (Fig. 6A) showed a combined peak (^2^H_9_-tCho’) of the overlapping resonances of deuterated Cho, PC, and GPC at 3.2 ppm, and a peak assigned to betaine at 2.25 ppm (^2^H_9_-tCho’ = ^2^H_9_-tCho - ^2^H_9_-betaine). Spectral fitting showed that tCho’ reflected 93.1 ± 3.6 % and betaine 6.9 ± 3.6 % of the [^2^H_9_-tCho] detected in vivo in the IV group (n = 8). In the PO group, tCho’ was 72.0 ± 8.2 % (p < 0.001 vs. IV) and betaine 28.0 ± 8.2 % (p < 0.001 vs. IV) of the [^2^H_9_-tCho] detected in vivo (n = 6).

**Fig. 6:**
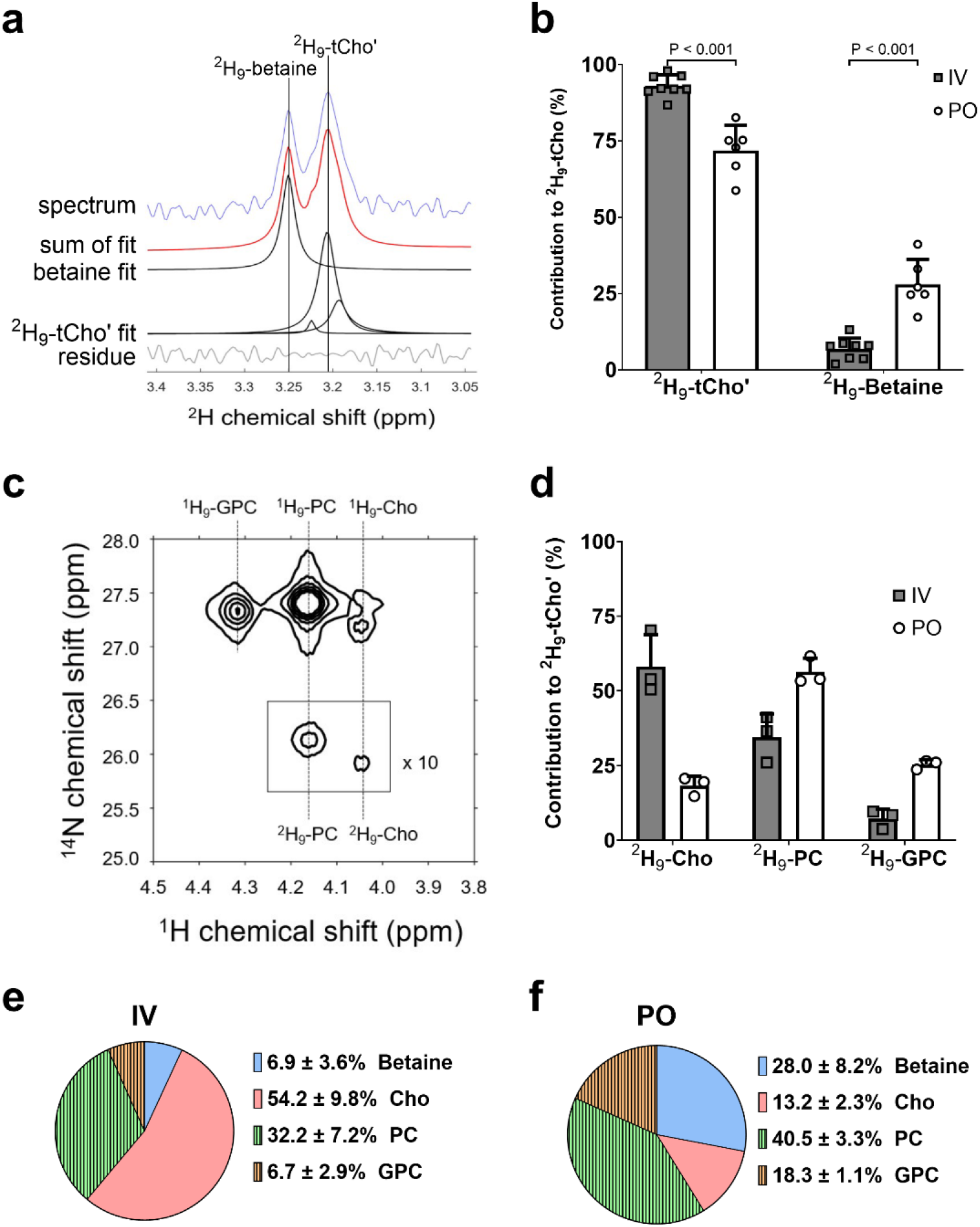
Ex vivo methods and quantification of individual ^2^H_9_-tCho metabolites. **a** Example of a ^2^H NMR spectrum acquired from a pooled tumor tissue sample of 2 animals following 3 days of PO-administered ^2^H_9_-Cho. Spectral fitting of the ^2^H_9_-tCho’ (combined peaks of ^2^H_9_-Cho, ^2^H_9_-PC and ^2^H_9_-GPC) and ^2^H_9_-betaine peaks is shown. **b** Relative contribution of ^2^H_9_-tCho’ and ^2^H_9_-betaine to ^2^H_9_-tCho. Individual points within bar graphs represent tumor samples from 2-3 animals combined, values are shown as mean ± SD for the IV group (n = 8) and the PO group (n = 5). All P values represent the results of two-tailed, independent sample, non-parametric t-test. **c** Example of a 2D ^1^H-^14^N HSQC NMR spectrum acquired from the same sample shown in (**a**). Note the peaks of both protonated and deuterated forms of the Cho metabolites. The intensity of deuterated metabolites was scaled 10-fold. **d** Relative contribution of ^2^H_9_-Cho, ^2^H_9_-PC, ^2^H_9_-GPC to ^2^H_9_-tCho’ for IV (n = 3) and PO (n = 3) administration groups. **e-f** Combined breakdown of the different molecules contributing for the *in vivo* [^2^H_9_-tCho] signal for the IV (**e**) and PO (**f**) groups. The shaded sections (^2^H_9_-PC and ^2^H_9_-GPC) in the pie chart indicate the contribution from active tumor metabolism.

To identify the different Cho species in the remaining ^2^H_9_-tCho’ peak, 2D ^1^H-^14^N 2D HSQC was performed, and the ratios of ^2^H_9_-Cho, ^2^H_9_-PC, and ^2^H_9_-GPC to ^2^H_9_-tCho’ were calculated. The corresponding values for the IV group (n = 3) were 58.2 ± 10.6% for ^2^H_9_-Cho, 34.6 ± 7.7% for ^2^H_9_-PC, and 7.2 ± 3.1% for ^2^H_9_-GPC, while for the PO group (n = 3), they were 18.3 ± 3.1% for ^2^H_9_-Cho, 56.3 ± 4.6% ^2^H_9_-PC, and 25.4 ± 1.6% ^2^H_9_-GPC. To account for the presence of betaine within the overall tCho signal in vivo, tCho’ metabolite fractions were multiplied by the tCho’ contribution from the ^2^H-NMR data. This enabled a comprehensive breakdown of the in vivo ^2^H_9_-tCho peak composition for both groups. In the IV group, the in vivo ^2^H_9_-tCho peak labeling consisted of 54.2 ± 9.8% Cho, 32.2 ± 7.2% PC, 6.7 ± 2.9% GPC, and 6.9 ± 3.6% betaine. For the PO group, the tCho signal breakdown was 13.2 ± 2.3% Cho, 40.5 ± 3.3% PC, 18.3 ± 1.1% GPC, and 28.0 ± 8.2% betaine (Fig. 6 e,f).

The 2D ^1^H-^14^N 2D HSQC data allowed quantification of both the protonated and deuterated forms of Cho, PC, and GPC, from which the fractional enrichment (FE) was calculated (Figure 6D). In the IV group (n = 3), FE was 0.303 ± 0.067 for Cho, 0.082 ± 0.021 for PC, and 0.029 ± 0.014 for GPC. FEs in the PO group (n = 3) were 0.081 ± 0.037 for Cho, 0.054 ± 0.017 for PC, and 0.048 ± 0.015 for GPC.

## Methods

### Brain Tumor Animal Model

All animal procedures were approved by the Yale University Institutional Animal Care and Use Committee. Brain tumors were induced in 26 Fischer 344 rats (Charles River, CT, USA) (235 ± 10 g) by intracerebral injection of 10,000 RG2 cells, as previously described ^15^. During the injection procedure, rats were continually anesthetized through inhalation of 2-3% isoflurane via a nose cone. Using aseptic techniques, a burr hole was made in the skull to inject a 5μL sterile saline solution with the RG2 cells. A Hamilton syringe (26 gauge) attached to a motorized injector and guided by a stereotaxic instrument was used to administer the cell solution at a rate of 1 μL/min. Peri- and post-operative care included the use of analgesics: Lidocaine at <7 mg/kg, Meloxicam at 1–2 mg/kg, and Buprenorphine at 0.01–0.05 mg/kg. After the tumor cell implantation, animals were monitored daily for recovery, and anatomical MRIs were performed regularly to monitor tumor growth.

### Choline Administration

For rats in the IV administration group (n = 14), the infusion protocol replicated the protocol conducted by Ip et al. (2023). In summary, catheters consisting of a 1-inch needle (30 gauge) and polyethylene tubing (PE10, Instech Laboratories, Plymouth Meeting, PA, USA) were placed in the rat’s lateral tail vein to allow for administration during the imaging studies. Twenty minutes before the Cho infusion, atropine sulfate (0.465 mg/kg, or 0.2 mg/kg free atropine base) was injected into the tail vein to counteract any cholinergic effects. ^2^H_9_-choline chloride (Cambridge Isotopes Laboratories, Cambridge, MA, USA) was dissolved in sterile water (400 mM) and administered through the catheter using a three-step bolus-continuous infusion protocol over 36 min, at a dose of 285 mg of the choline base per kg body weight. This infusion protocol resulted in an infused volume of 6.3 mL per kg body weight.

For rats in the PO administration group (n = 12), oral gavage was performed with flexible polypropylene feeding tubes to administer a solution of ^2^H_9_-choline chloride dissolved in sterile water. The protocol resulted in an administered dose of 50 mg/kg of the choline base daily for three consecutive days. This dose was chosen based on the established choline daily upper limit for human adults set at 3500 mg and an assumption of average human adult body weight of 70 kg ^20^.

### In Vivo DMI and MRI Imaging

During imaging, all animals were free-breathing, anesthetized with isoflurane using ~60% O_2_ and ~40% N_2_O as carrier gas mixture, delivered through a nosecone ^15^. A heating pad was used to maintain body temperature at ~37° C.

All animals in the IV administration group started the in vivo scanning process concurrent with the start of ^2^H_9_-choline chloride IV infusion. All animals in the PO group were scanned after completion of the oral gavage on day 3 of administration (~2 hours after the gavage). Two animals were scanned ~2 hours following oral administration on days 1 and 2 to follow any buildup of ^2^H_9_-tCho over time.

All imaging studies were performed on an 11.74 T magnet interfaced to a Bruker Avance III HD spectrometer running on ParaVision 6.01, using a 20 mm × 15 mm elliptical ^2^H surface coil. Two additional orthogonal 20 mm ^1^H surface coils driven in quadrature were used for anatomical imaging and B_0_ shimming. Following scout MRIs to check animal position, CE T_1_W MRIs were acquired using a multi-slice spin-echo pulse sequence, repetition time (TR) of 1,000 ms, and echo time of 6.4 ms, 10-20 min after a 150–200 μL bolus of the T_1_ contrast agent gadopentetate dimeglumine (Magnevist®, Bayer, NJ, USA) was administered subcutaneously. Following the acquisition of the anatomical MRIs, 3-D B_0_ mapping and shimming were performed. After shimming (using second-order spherical harmonics), the FWHM linewidth of water was 25–35 Hz in a ~350 μL volume. Deuterium MR signal acquisition used a pulse-acquire sequence extended with 3D spherical phase-encoding during the initial 0.6 ms after excitation. DMI data was acquired as an 11 × 11 × 11 matrix in a 27.5 mm × 27.5 mm × 27.5 mm field of view (TR = 400 ms, 8 averages) and using spherical k-space sampling, resulting in a 2.5 × 2.5 × 2.5 mm = 15.6 μL nominal spatial resolution for a 36 min total scan time.

### Collection and Processing of Brain and Tumor Tissue Samples

Immediately following the completion of the final DMI scan, animals were euthanized by isoflurane overdose and decapitated. Tumor tissue (47.3 - 173.1 mg) was harvested, frozen in liquid N_2,_ and stored at –80 °C until further processing. Tissue metabolite samples were homogenized using a bead mill (Omni International, Kennesaw, GA, USA) and a standard methanol-HCl extraction protocol ^24^. For ^2^H NMR analysis, samples were dissolved in 600 μL of a water-based 100 mM phosphate buffer, pH = 7.3, using 5 mm NMR tubes. After the ^2^H NMR scans, the samples were dried, and groups of 2-3 samples were recombined during resuspension to achieve a total sample mass of ~200 mg, to enhance the detection sensitivity for 2D ^1^H-^14^N HSQC NMR. The solvent used for 2D ^1^H-^14^N HSQC NMR was 300 μL D_2_O-based pH-buffered solution, which was transferred to a Shigemi NMR tube, susceptibility matched to D_2_O (ATS Life Sciences, Wilmad, NJ, USA).

### Ex-Vivo NMR Spectroscopy

High-resolution ^2^H NMR and 2D ^1^H-^14^N HSQC NMR scans of tumor tissue metabolite extracts were performed on a 500 MHz Bruker Avance MR spectrometer (Bruker Instruments, Billerica, MA, USA) using the ^2^H channel typically used for signal locking. Deuterium NMR experiments were performed at 76.77 MHz, using a standard pulse-acquire sequence (TR: 1.5 s, ns: 2048 - 7,200), with locking disabled. 2D ^1^H-^14^N HSQC NMR spectra were acquired as 2048 complex points over a 5 kHz spectral width for ^1^H and 128 t_1_ increments over a 0.4 kHz spectral width for ^14^N, as described in de Graaf et al. ^25^. All experiments were performed at 298 K.

### Data Processing

All DMI data were processed in MATLAB (The Mathworks, Natick, MA, USA). DMIWizard, an in-house graphical interface within MATLAB, was used to perform the spectral fitting of the ^2^H MRSI spectra. Fitting was accomplished through a linear combination of two model spectra: the natural abundance water and the combined tCho peak. Spectral fitting included a variable zero-order and fixed first-order phase correction. Upon successful fitting, the tCho peak was quantified by converting signal amplitude to concentration values using the water signal as an internal concentration standard. We estimate the HDO internal concentration standard to be 13.2 mM, assuming a water content of 80% for both NAB and tumor tissue, and 0.015% ^2^H natural abundance ^26^. Due to the similar T1 values for water and tCho, no corrections for T1 relaxation were performed ^7,15,27^. The resulting concentrations of ^2^H_9_-tCho were used to create metabolic maps of the brain and to quantify the average concentration of ^2^H_9_-tCho within tumors and NAB.

To create the metabolic maps, 2D interpolation was used by convolving the nominal DMI data with a Gaussian kernel, whereby the convolution provided an inherent Gaussian smoothing of 1.2 - 1.8-pixel widths, as previously described ^7^. For figures, the interpolated DMI maps displayed as amplitude color maps were overlaid on anatomical MRI.

The tumor-specific ^2^H_9_-tCho concentration was calculated by averaging the [^2^H_9_-tCho] values from all the tumor-containing MRI voxels. Tumor and contralateral NAB VOIs were generated via manual segmentation on CE T_1_W MRIs in ITK-SNAP (Version 3.8.0, www.itksnap.org)^28^. The resulting segmentation matrices were multiplied by DMI data using an in-house written script in MATLAB. The script extrapolated the 11 × 11 × 11 DMI matrix to match the 88 × 88 × 88 MRI matrix, resulting in each DMI voxel containing 512 MRI voxels (8 × 8 × 8). Each MRI-based voxel was assigned the [^2^H_9_-tCho] of the overlapping DMI voxel (Supplemental File, Fig. S1).

^2^H NMR data were processed in NMRWizard, an in-house produced graphical interface within MATLAB. NMRWizard was used to perform spectral fitting of the ^2^H MRS spectra. Peak fitting included a variable zero-order and fixed first-order phase correction and was accomplished through a linear combination of four model spectra for the trimethyl peaks of Cho (3.18 ppm), PC (3.20 ppm), GPC (3.22 ppm) and betaine (3.25 ppm). The betaine peak showed minimal overlap with other peaks but because the Cho, PC, and GPC peaks were overlapping, their amplitude estimates were combined into a single peak labeled as tCho’. To resolve the contribution of the different metabolites within the tCho’ peak, 2D ^1^H-^14^N HSQC NMR was performed.

The data from 2D ^1^H-^14^N HSQC NMR were processed following the procedures outlined in de Graaf et al., 2025 (De Graaf et al., 2025). In short, data were processed offline in NMRWizard where spectra were apodized (–1 Hz exponential, 5 Hz Gaussian in both dimensions), zero-filled to 4K x 4K data points, Fourier transformed, and displayed in absolute value. The resulting 2D spectra showed six trimethyl peaks, corresponding to the deuterated and non-deuterated forms of Cho, PC, and GPC, respectively. Non-deuterated species were quantified with a 2D least-squares fitting using a linear combination of model spectra. Before the quantification of deuterated species, a Hankel singular value decomposition (HSVD) algorithm was applied to remove the signal of the non-deuterated species ^29^. This was performed to prevent the much higher signal from the non-deuterated species from bleeding into the relatively smaller peaks of the deuterated species. Following the HSVD application, deuterated species were also quantified with the same peak fitting procedure as the non-deuterated species.

### Statistics and Data Analysis

All statistical tests were performed using GraphPad Prism version 10.4.0 for Windows (Boston, MA, USA, www.graphpad.com). A Shapiro-Wilk test performed on in vivo [^2^H_9_-tCho] data confirmed the normal distribution of data from tumor in the IV (n = 10, p = 0.18) and PO (n = 9, p = 0.34) groups, as well as NAB in the IV (n = 10, p = 0.46) and PO (n = 9, p = 0.14) groups. Two-tailed, independent samples t-tests were used to compare in vivo [^2^H_9_-tCho] values between tumor and NAB, as well as between IV and PO groups. Tumor-to-NAB was analyzed as the ratio of [^2^H_9_-tCho] in the tumor over the [^2^H_9_-tCho] in NAB for each animal. The analysis of high-resolution ^2^H NMR data included calculating betaine/(betaine+tCho’) to find the contribution of betaine to the tCho signal. The tCho’ fraction of tCho (tCho’/[betaine+tCho’]) was used to determine the contribution of Cho, GPC, and PC (obtained from the 2D NMR) to the tCho signal. A two-tailed, independent samples t-test was used to compare betaine and tCho’ between the IV and PO groups. Data from 2D HSQC NMR and resulting fractional ^2^H enrichment values were analyzed using descriptive statistics because of small sample sizes (n = 3). Fractional ^2^H enrichment was calculated as ^2^H_9_-X / (^1^H_9_-X + ^2^H_9_-X), where X represents Cho, GPC, or PC.

## Discussion

In the ongoing search for a metabolic imaging technique that can provide high image contrast between tumor and NAB, while also being cancer-specific, we investigated the performance of DMI with orally administered ^2^H_9_-Cho. The involvement of Cho in the synthesis of phospholipids via the Kennedy pathway makes Cho uptake and metabolism an excellent target for cancer-specific imaging. In this study, we built on previous work that showed high image contrast in rat brain tumor models both during and following IV infusion of a relatively high dose of deuterated Cho. We demonstrated that even when using a significantly lower dose, equivalent to the maximum recommended daily Cho intake as a nutritional supplement for adults, this protocol resulted in significant image contrast in a rodent model of GBM. This result confirms our hypothesis, which was based on the previous work that showed a significant ^2^H_9_-tCho peak 24 hours after the initial IV infusion of ^2^H_9_-Cho. The high level of labeling with low oral doses can seem counterintuitive at first, but can be explained by the ‘trapping’ of ^2^H-label from ^2^H_9_-Cho in its immediate downstream metabolite PC, and to a lesser extent in GPC. Therefore, consecutive oral doses can build up the labeled PC pool, which turns over at a slow rate compared to the phosphorylation of Cho by CKA. This mechanism is comparable to using the radioactively labeled glucose analog, ^18^F-2- deoxyglucose (FDG), as a tracer for positron emission tomography (PET). The lack of the glucose hydroxyl group on the 2-carbon, replaced by ^18^F through chemical modification, prevents further metabolism of 2-deoxyglucose, and leads to the accumulation of FDG ^30^. In contrast, within the time span of our experiment, the deuterium label of ^2^H_9_-Cho is ‘trapped’ naturally in PC. The rapid conversion of free Cho to PC and relatively slow further conversion was also previously observed in vitro, in breast cancer cells ^31^. The ability to accumulate ^2^H label in PC is key to using a low dose of ^2^H_9_-Cho while still achieving significant SNR and tumor-to-NAB image contrast.

Although the tCho DMI maps were similar for the two dosing strategies, the make-up of the ^2^H_9_- tCho peaks was not equal. In the IV group, a significant portion of tumor ^2^H_9_-tCho originated from free Cho, which can be intra- or extracellular in the tumor. In contrast, in the PO group, the free Cho contribution was relatively low, and the bulk of the ^2^H_9_-tCho peak consisted of labeled PC and GPC. Both the sum and each individual contribution of labeled PC and GPC to the ^2^H_9_-tCho peak in the PO group were larger than in the IV group. Because PC and GPC are the result of activity by CKA and the Kennedy pathway, the 3-day PO protocol led to tCho-DMI maps that were arguably more specific for cancer metabolism as compared to the IV approach. In the IV group, the bulk of the ^2^H_9_-tCho peak originated from free Cho, which could be located in the blood, extracellularly, or in the intracellular space, and is therefore not representative of tumor metabolism.

Oral administration of ^2^H_9_-Cho also results in the formation of deuterated betaine, which contributes to the ^2^H_9_-tCho peak and the tumor-to-NAB image contrast. Betaine is irreversibly synthesized from Cho predominantly in liver tissue, and appears to be easily transported into the tumor. We could not determine whether this tumor betaine resided intracellularly or extracellularly. In either scenario, we consider betaine a non-specific contributor to the image contrast because, to the best of our knowledge, conversion of betaine to Cho has only been described in fungal cells ^32^.

We chose 3 days for the oral loading regimen because by the third day, the pilot studies showed in vivo SNR and image contrast for ^2^H_9_-tCho that was comparable to the single IV studies. It is currently unclear whether a longer period of oral loading would result in yet higher SNR of ^2^H_9_-tCho. Based on the relatively low fractional ^2^H enrichment (<10%) of ^2^H_9_-PC and GPC, there appears to be significant room to increase the ^2^H-labeling over a longer period of time. This potential higher SNR could be traded for higher spatial resolution of the DMI-based metabolic maps or shorter scan times. To what extent betaine would also be increased and contribute to the in vivo ^2^H_9_-tCho peak is unknown; therefore, future experiments to optimize the oral loading period will need to include analyses of the different Cho tumor-specific and non-specific metabolites.

From previous work, we learned that the in vivo ^2^H_9_-tCho peak amplitude was ~25% lower the day after a high-dose IV infusion ^15^. Currently, it is unclear how long it takes for the in vivo ^2^H_9_-tCho peak to disappear after 3 days of oral dosing. This becomes particularly relevant if tCho-DMI is explored as an early biomarker of treatment response. The expectation would be that a treatment-responsive tumor has lower or no phospholipid synthesis activity and thus reduced demand for Cho metabolism, which would result in low levels of ^2^H_9_-tCho after oral dosing with deuterated Cho. For tCho-DMI to be useful in a comparison between pre- and post-treatment, the ^2^H_9_-tCho from the pre-treatment scan would ideally have vanished before both the start of treatment and the follow-up tCho-DMI scan.

The tCho-DMI-based metabolic image contrast in vivo between tumor and NAB was high, in part because of the undetectable ^2^H_9_-tCho peak in NAB. The reported measurements of [^2^H_9_-tCho] in normal brain are in fact an overestimation as a result of peak fitting of the random noise peaks in the 2.2 ppm area of the spectrum, since we did not visually observe a ^2^H_9_-tCho peak in NAB. While normal brain does require Cho for the synthesis of the neurotransmitter acetylcholine, uptake was below the in vivo detection limit of our 36 min tCho-DMI scan ^9^. This observation highlights the potential role of the leaky blood-brain barrier that is typical for GBM, in the uptake of the blood-borne Cho. The regions with high ^2^H_9_-tCho levels also spatially match the areas with high contrast on CE T_1_W MRI, the indicator of the compromised blood-brain barrier. However, the spatial resolution of tCho-DMI is much lower than of the anatomical MRI, and therefore, small differences in area and location cannot be observed. To better understand the role of a leaky blood-brain barrier, animal models of low-grade brain tumors that have an intact blood-brain barrier need to be studied.

The RG2 model of GBM harbors a number of features observed in human GBM ^33^. Yet, this model is biologically not the best representation of the human disease. For example, unlike human brain tumors, RG2 rat tumors are relatively homogenous ^34^. However, the goal of this work is not to discover new cancer biology, but to visualize well-established aspects of brain tumor metabolism and create high tumor-to- NAB image contrast. For this purpose, tumors grown in F344 rat brain following injection of the isogeneic RG2 cells are an appropriate animal model.

Currently, there is limited insight into whether DMI combined with ^2^H_9_-Cho administration would be beneficial in other tumor types as well. Veltien et al. have shown high uptake of ^2^H_9_-Cho during IV infusion in a mouse model of renal cell carcinoma ^35^. However, that study was focused on demonstrating that DMI data can be acquired during the co-infusion of both ^2^H_9_-Cho and [6,6’-^2^H_2_]-glucose, informing on two distinct metabolic pathways in a single imaging study. The study was performed in tumors growing subcutaneously, and therefore, we cannot speculate on any image contrast with surrounding normal kidney tissue. An earlier study in a mouse breast cancer model also shows an example of high ^2^H_9_-tCho signal after IV infusion of deuterated Cho ^31^. But the nature of rodent breast cancer models also does not facilitate comparison with non-tumor tissue, and probably needs to be performed in humans to get a clear answer.

Translation of tCho-DMI includes both the DMI data acquisition and the ^2^H_9_-Cho administration. DMI acquisition in combination with oral [6,6’-^2^H_2_]-glucose intake is currently being used in human subjects by different research groups and at different magnetic field strengths and is thus proven to be translatable ^36–39^. IV infusion of deuterated Cho would not be very challenging in a clinical setting, but might require the co-infusion of atropine to counter possible side effects. In contrast, an oral dosing regimen of ^2^H_9_-Cho at the dose used in this study would not need atropine and have other benefits. Unlike IV contrast agents, orally administered choline potentially reduces the need for clinical staff and equipment for IV administration. A clinical version of the PO administration could rely on ingestion of tablets of ^2^H_9_-Cho for several days leading up to a DMI scan. Initial first-in-human tCho-DMI studies would need to focus on dose, loading period, SNR, and tumor-to-NAB image contrast. Further, plasma and excised tissue analysis would be needed to fully understand the dynamics driving the labeling in a brain tumor. The different metabolites contributing to the tCho peak would need to be defined using ex vivo high-resolution NMR, as was done in this study. Because most high-grade brain tumors are resected as part of standard-of-care, collecting tumor tissue after ^2^H-labeled Cho ingestion should be feasible to achieve. A clinical version of tCho-DMI could also include oral intake of [6,6’-^2^H_2_]-glucose, 60 to 90 minutes before the DMI scan. As shown by Veltien et al., the ^2^H_9_-tCho peak does not overlap with peaks of glucose or its metabolites, at least in a preclinical setting at ultra-high magnetic field ^35^. Based on ^2^HMR spectra acquired in human brain at 4T after [6,6’-^2^H_2_]-glucose intake, it can be expected that the ^2^H_9_-tCho peak would minimally overlap with the labeled glutamate+glutamine peak, even at a lower magnetic field strength ^7,38^. This indicates that at clinical field strengths, performing DMI combined with both [6,6’-^2^H_2_]- glucose and ^2^H_9_-Cho administration would be feasible as a single scan. Lastly, recent developments that make it feasible to acquire both DMI and MRI in parallel have resulted in a comprehensive anatomical and metabolic neuro-imaging protocol. Being able to acquire both the anatomical and metabolic imaging data without requiring extra scan time is an important step towards clinical integration ^40,41^.

This study demonstrated that serial oral administration of ^2^H_9_-Cho can achieve image contrast in DMI-based metabolic maps comparable to IV administration, while using a dose recommended for human adults. The predominant contributions of Cho metabolites to the maps implied that the image contrast is not merely caused by Cho uptake but predominantly driven by tumor Cho metabolism. This approach offers a noninvasive, clinically translatable alternative for metabolic imaging of brain tumors, providing a foundation for future studies integrating DMI techniques into clinical practice.

## List of Abbreviations

DMI: Deuterium metabolic imaging
MRSI: Magnetic resonance spectroscopic imaging
tCho: total choline (= choline, phosphocholine, glycerophosphocholine, and betaine); tCho’total choline excluding betaine
HSQC: Heteronuclear single quantum coherence
Cho: Choline
PC: Phosphocholine
GPC: Glycerophosphocholine,
GBM: Glioblastoma
PO: Oral administration
IV: Intravenous
CE: T_1_W
MRI: Contrast enhanced T_1_ weighted magnetic resonance imaging
CKA: Choline kinase alpha
NAB: normal- appearing brain
VOI: Volume of interest
SNR: signal-to-noise ratio
TR: Repetition time
FWHM: Full width at half maximum
FE: fractional enrichment.

