## Supplemental File for "Oral Intake of Deuterated Choline at Clinical Dose for Metabolic Imaging of Brain Tumors"

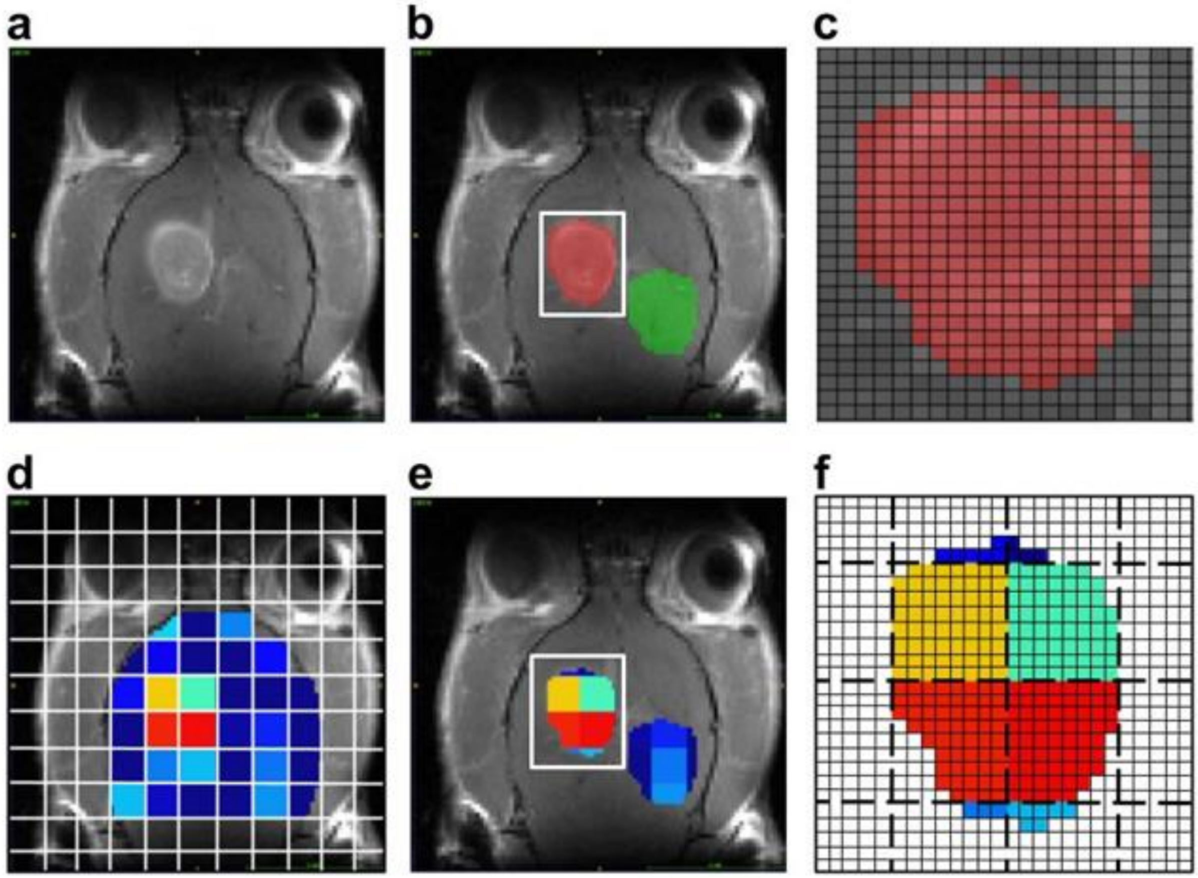


**Figure S1: Segmentation and data averaging of brain areas.**

**a** Coronal slice of CE $T_1$W MRI from an RG2-bearing rat used to segment the volume of interest (VOI) for the tumor lesion. **b** Example VOIs representing tumor (red) and NAB (green) in the contralateral hemisphere. **c** Zoomed-in section of panel **b** (white square) showing the resolution of the MRI (black grid, based on 88 x 88 x 88 matrix) used for segmentation. **d** Color-coded $^2$H MRSI voxels within the relevant brain area, overlaid on anatomical MRI. The white grid illustrates the resolution of the DMI acquisition (white grid, based on 11 x 11 x 11 matrix). **e** Results of multiplying VOIs by $^2$H MRSI. **f** Zoomed-in section from panel **e** (white square). 2-D representation of the MRI matrix (solid lines) and $^2$H MRSI matrix (dashed lines). A total of 512 MRI voxels (8 x 8 x 8) fit into each $^2$H MRSI voxel. Color-coding reflects $^2$H MRSI-based quantification of [$^2H_9$-tCho]; white pixels = 0.
